# Microglia respond to elevated intraocular pressure and synapse loss in the visual thalamus in a mouse model of glaucoma

**DOI:** 10.1101/2024.10.25.619907

**Authors:** Jennifer L. Thompson, Shaylah McCool, Jennie C. Smith, Victoria Schaal, Gurudutt Pendyala, Sowmya Yelamanchili, Matthew J. Van Hook

## Abstract

Microglia are resident immune cells of the central nervous system and mediate a broad array of adaptations and responses during disease, injury, and development. Typically, microglia morphology is understood to provide a window into their functional state. However, it is apparent that they have the capacity to adopt a broad spectrum of functional phenotypes characterized by numerous morphological profiles and associated gene expression profiles. Glaucoma, which leads to blindness from retinal ganglion cell (RGC) degeneration, is commonly associated with elevated intraocular pressure and has been shown to trigger microglia responses within the retinal layers, at the optic nerve head, and in retinal projection targets in the brain. The goal of this study was to determine the relationship of microglia morphology to intraocular pressure and the loss of retinal ganglion cell output synapses in the dorsolateral geniculate nucleus (dLGN), a RGC projection target in the thalamus that conveys information to the primary visual cortex. We accomplished this by analyzing dLGN microglia morphologies in histological sections from DBA/2J mice, which develop a form of inherited glaucoma. Microglia morphology was analyzed using skeletonized Iba1-fluorescence images and fractal analyses of individually reconstructed microglia cells. We found that microglia adopted more simplified morphologies, characterized by fewer endpoints and less total process length per microglia cell. There was an age-dependent shift in microglia morphology in tissue from control mice (DBA/2J^Gpnmb+^) that was accelerated in DBA/2J mice. Microglia morphological measurements correlated with cumulative intraocular pressure, immunofluorescence labeling for the complement protein C1q, and density of vGlut2-labeled RGC axon terminals. Additionally, fractal analysis revealed a clear distinction between control and glaucoma dLGN, with microglia from ocular hypertensive DBA/2J dLGN tissue showing an elongated rod-like morphology. RNA-sequencing of dLGN tissue samples showed an upregulation of immune system-related gene expression and several specific genes associated with microglia activation and potential neuroprotective functions. These results suggest that microglia in the dLGN alter their physiology to respond to RGC degeneration in glaucoma, potentially contributing to CNS adaptations to neurodegenerative vision loss.

## Introduction

Glaucoma is an age-related neurodegenerative disorder of the visual system and one of the leading causes of blindness and visual impairment worldwide [1–4]. All forms of glaucoma are characterized by optic nerve atrophy and irreversible vision loss due to the progressive death of retinal ganglion cells (RGCs), which are the output neurons of the retina [5–7]. The two most prominent glaucoma risk factors are increasing age and elevated intraocular pressure (IOP). Since only IOP is modifiable, IOP-reduction forms the core of glaucoma management strategies and IOP-monitoring plays a pivotal role in identifying glaucoma suspects and predicting future progression [5]. Understanding the links between IOP and pathological changes in glaucoma is critical to establishing a full picture of disease progression and identifying potential diagnostic and therapeutic strategies.

Elevated IOP damages RGCs at the optic nerve head, where their axons converge before exiting the eye through a meshwork of glial and structural support cells and making their way to >50 retinal projection targets in the brain [8–10]. Within the brain itself, downstream circuit impairment is evidenced by glaucoma-specific dysfunction in multiple white matter tracts related to visual processing, and gray matter reductions resulting from transneuronal degeneration are apparent in retinorecipient nuclei and the visual cortices of glaucoma patients [10–16]. Evidence from experimental animal models suggests that some of the earliest afflictions in glaucoma may occur in the brain [17–23]. In mice with glaucoma for instance, RGC axon terminals in the superior colliculus show early signs of degeneration including bouton atrophy and mitochondrial abnormalities [24]. We have previously reported an IOP-related loss of RGC axon terminals and diminished RGC synaptic function along with atrophy and hyperexcitability of post-synaptic neurons in the dorsolateral geniculate nucleus (dLGN), a critical RGC projection target for conscious vision and the rodent analog to the primate lateral geniculate nucleus (LGN) [8,17,19,25,26].

Microglia are innate immune cells of the brain that perform essential tasks related to development, homeostasis, and maintenance of local circuits [27]. Microglia are also first-responders to tissue insult and injury, and their reaction is often characterized by a morphological transition towards an ameboid state and the adoption of a proinflammatory cytokine profile [27–29]. A systematic review of post-mortem tissue samples from human glaucoma patients concluded that amoeboid microglia are frequently observed near the optic nerve head (ONH) and are positive for markers of pro-inflammatory cytokines [30]. Microglia are also implicated as proinflammatory mediators of the biological response to IOP elevation for a variety of animal models [30–34]. The goal of this study was to determine whether and how elevated IOP and glaucoma impact microglia in the dLGN. Using DBA/2J mice, which develop elevated IOP (ocular hypertension, OHT) and glaucoma due to mutations in glycosylated protein nmb (*Gpnmb*) and tyrosinase-related protein 1 (*Tyrp1b*) genes [35–38], we find that dLGN microglia become less morphologically complex and adopt a bipolar or “rod-like” morphology [39–42] in a manner that correlated with IOP and loss of immunofluorescent staining for retinal ganglion cell axon terminals. We also found that microglia morphology changes correspond to increased labeling of complement C1q in the dLGN, potentially involved in targeting degenerating RGC axon terminals and other debris for microglial phagocytosis [27,43,44].

Finally, bulk RNA sequencing of dLGN pointed toward increased expression of genes associated with immune responses.

## Materials and Methods

### Animals

All protocols involving animals were approved by the Institutional Animal Care and Use Committee at the University of Nebraska Medical Center. Male and female DBA/2J (Jackson Labs #000671, RRID:IMSR_JAX:000671) and DBA/2J^Gpnmb+^ (Jackson Labs #007048, RRID:IMSR_JAX:007048) mice were bred onsite and housed socially under a 12/12-hour light/dark cycle with free access to food and water. Eye pressures were measured (in mmHg) using a handheld rebound tonometer (TonoLab, iCare) calibrated for mouse eyes. Baseline IOPs were obtained when mice were approximately 3 months-old, then data collection continued monthly for the duration of the study. All subsequent recordings were performed at or near the same time of day to minimize the effects of diurnal variance. To avoid stress-induced IOP elevations, measurements were obtained while mice were under light anesthesia (2% isoflurane) within five minutes of losing their righting reflex. Because six consecutive, low-error measurements from the central cornea are required to generate a single readout, a total of 18 measurements were obtained per eye per timepoint, and each IOP value represents the average of three readouts.

### Immunofluorescence staining

At 4-, 9-, and 12-months of age, mice of each strain were euthanized via CO_2_-inhalation followed by cervical dislocation. Immediately afterwards, brains were dissected, rinsed, and drop-fixed in 4% PFA (in PBS: phosphate buffered saline) for 4hr, shaking gently at room temperature. For cryopreservation, after three 10-minute washes, brains were immersed in a 30% sucrose solution and kept at 4°C until saturated. Afterwards, brains were placed into tissue molds and embedded in 3% agar. 50µm coronal sections containing the dLGN were obtained using a vibratome (Leica VT1000S), then mounted onto SuperFrost Plus slides and stored at -20°C until use. No animals or tissues were excluded.

For immunostaining, DBA/2J and DBA/2J^Gpnmb+^ tissue sections containing the dLGN core were used. After a PBS rinse, slides were blocked and permeabilized for 1hr at room temperature in a 7.4 pH buffer containing 5% donkey serum, 5% goat serum, and 0.5% Triton X-100. To visualize RGC axon terminals, an antibody against vesicular glutamate transporter 2 (anti-VGluT2; Cat#6D2092; Millipore, RRID: AB_2665454) was used at a concentration of 1:250, while microglia were identified using an antibody against ionized calcium binding adaptor protein 1 (anti-Iba1; Cat#019-19741, Wako; RRID: AB_8395054) at a concentration of 1:500. Together, primary antibodies were diluted in fresh buffer solution and slides were stained over the weekend (3 overnights) in order to enhance the resolution of finer microglial processes. To identify C1q dLGN coronal sections from a subset of DBA/2J (n=32) and DBA/2J^Gpnmb+^ mice (n=25 mice were similarly blocked and permeabilized before overnight incubation in buffer diluting a knockout validated anti-C1q antibody (Cat#ab182451, Abcam; RRID: AB_2732849) at 1:500. Primary incubations were followed by a series of six ten-minute washes and an additional 1hr block. For visualization, slides were incubated in the dark for 3h at room temperature in buffer containing the appropriate secondary antibodies—donkey anti-rabbit IgG (H+L), Alexa Fluor 568-conjugated (Cat#A10042; RRID: AB_2534017) and/or goat anti-guinea pig IgG (H+L), Alexa Fluor 488-conjugated (Cat#A-11073; RRID: AB_2534117)— each diluted 1:200. Unbound secondary antibodies were cleared (3x10min washes) and slides were rinsed in dH_2_O prior to being coverslipped with VECTASHIELD HardSet (Cat#H1400).

For fluorescently labeled DBA/2J and DBA/2J^Gpnmb+^ tissues, two photon microscopy was used to acquire a series of image stacks (1 µm Z spacing; 370 x 370 µm; 1024 x 1024px) through the central dLGN core. A single image plane was used to generate a spatially-resolute field of fluorescent puncta corresponding to the vGluT2-bound signal. A density measurement was then generated via incorporating the image acquisition area into the particle analysis output. For measuring C1q Intensity, Z-stacks were compressed by group-averaging with a Z-projection method, then backgrounds were subtracted locally using a rolling ball radius set to 50 pixels.

Then the average pixel intensity values were determined for each frame and the maximum average intensity value was identified for each sample.

### Global Skeleton Analysis

Skeleton analysis of microglia was performed using a method adapted from Young and Morrison (2018) [45]. To achieve a uniform thickness across samples, slices in the Iba1+ channel were trimmed to span a thickness of 40µm. Slices were then projected onto a single plane based on maximum intensity values to generate a two-dimensional surface that enhanced the signal arising from fine processes while maintaining the morphological integrity of Iba1+ cells. This was converted to an 8-bit black and white image. Edges were enhanced with an unsharp mask filter set to a 0.5px sigma radius and a 0.6 mask weight. Impulse noises were removed using the *Despeckle* function before and after a global threshold (22%) was applied using the default method (modified IsoData algorithm). A binary close command was run to bridge gaps in adjacent microglial processes that had perhaps eroded or provided weak signal. Then dark noise was smoothed by running the *Remove Outliers* command set to a 7px radius and a deviation threshold of 50 against the surrounding median. Final skeletonized images were generated by running the *Skeletonize (2D/3D)* command, and analyzed with the *Analyze Skeletonize 2D/3D* plugin. From the output windows, endpoint and branch length data were manually trimmed to exclude small (<1µm) incomplete (<2 endpoints) branches. To calculate global averages per microglia, Iba1+ cell n-counts were performed by using the *Cell Counter* plugin and scrolling through the 40µm Z-stacks.

#### Fractal Analysis

To analyze morphology of individual microglia [45–47], 216 Iba1+ cells were hand-traced throughout the same set of 40µm image stacks. For randomization, one Iba1+ cell in the central region of each quadrant (4 per dLGN) was selected on the basis of being fully contained within the tissue volume. A 3D Gaussian Blur filter was applied with a 1.0 sigma radius on each plane. In a duplicate set, images were converted to masks with the “redirect to” option of the *Analyze Particles* function. Selected Iba1+ cells were traced through the masked volume using the *Blow and Lasso Segmentation Tool*, and ROIs were frequently redirected to the unmasked, filtered z-stacks for reference. Final 2D outlines were added to the ROI Manager and saved in sets of four. One-by-one, each trace was converted into its own mask, resulting in a set of two-dimensional binary images each containing a single microglia silhouette. ROIs were outlined using the *Binary* options and whole images were analyzed using the *FracLac* plugin [47] set to the (box-counting method; 4 start positions; 45% largest grid size) with *Hull & Circle* results selected.

Thresholds for masking were determined on a sample-by-sample basis, and occasionally single pixels were manually inserted for the purpose of filling or recreating gaps (i.e. between pseudo-overlapping processes). All final outlines were meticulously retouched to align with the true, unmasked signal.

### RNA sequencing

Bulk RNA sequencing was performed with a total of eight samples from 9-month-old DBA/2J and DBA/2J^Gpnmb+^ control mice. To isolate dLGN tissue, brains were dissected into an RNAse-free slush of phosphate buffered saline and cut into 500 micron-thick sections on a vibratome. The dLGN from both hemispheres was then identified and dissected free. dLGN tissue from three mice was pooled into each sample and weighed before being flash frozen on dry ice. Total RNA was isolated from dLGN tissue using the Direct-Zol RNA kit (Zymo Research, Irvine, CA). Approximately 1 µg of RNA per sample was sent on dry ice to LC Sciences for RNA sequencing with their Poly(A) RNA sequencing service. The Poly(A) RNA sequencing library was prepared using the TruSeq-stranded-mRNA protocol (Illumina) and purified using oligo-(dT) magnetic beads. After purification and DNA library construction, quality control and quantification was performed using the 2100 Bioanalyzer High Sensitivity DNA Chip (Agilent Technologies) and paired-ended sequencing performed using the NovaSeq 6000 sequencing system (Illumina). Cutadapt and Perl scripts were used by LC Sciences for bioinformatics analyses to remove reads containing adaptor contamination, low-quality bases, and undetermined bases and sequence quality assessed using FastQC. HISAT2 was used to map reads to the genome and reads of each sample were assembled using StringTie, after which transcriptomes were merged to reconstruct a comprehensive transcriptome. StringTie and Ballgown were then used to estimate the expression levels of all transcripts.

### Statistics

All statistical tests were performed in GraphPad Prism 10 unless noted otherwise. To represent the cumulative effect of IOP across time, longitudinal IOP data were compressed into a single AUC integral (area under the curve in mmHg*days). Two-way ANOVAs, factoring strain and age, were used for group comparisons, followed by Šídák-corrected pairwise comparisons. Comparisons involving individual Iba1+ features were performed using nested oen-way ANOVAs followed by Tukey’s HSD tests. To test explanatory relationships between variables, simple linear regressions were used and follow-up slope comparisons (two-sided) were performed. P-values < 0.05 were considered statistically significant. Data are reported as mean ± standard deviation in bar graphs and 95% confidence intervals in regression plots.

## Results

To determine the extent and time-course of OHT in our DBA/2J colony we recorded IOP on a monthly basis from the left and right eyes of 39 DBA/2J mice and 36 DBA/2J^Gpnmb+^ controls using a handheld rebound tonometer (Fig.1A-B), then used left-eye data to compare IOP averages between strains at three ages relevant to the onset and severity of DBA/2J glaucoma (4-, 9-, and 12-months, Fig. 1C). As expected, a two-way ANOVA testing the effects of age and strain on final IOP values resulted in a significant interaction term [*F*(2,69)=6.35, *p*=0.003] due to age-related OHT being relatively exclusive to older DBA/2J eyes. Although DBA/2J^Gpnmb+^ mice carry the *Tyrp1^b^* allele that partakes in mediating DBA/2J glaucoma, without the combinatory *Gpnmb^R^*^150*X*^ mutation, they are spared from severe iris disease and do not become ocular hypertensive [36,37]. In line with this, DBA/2J^Gpnmb+^ IOPs were indistinguishable by age (4mo-9mo: *p=0.75*; 9mo-12mo: *p=0.99*; Tukey’s multiple comparisons), with IOP averages of 11.6±1.4 mmHg at 4-months, 12.8±1.9 mmHg at 9-months, and 13.0±2.4 at 12-months (Fig.1C). In contrast, age-related IOP elevations are clearly apparent in DBA/2J eyes in both 9- and 12-month conditions (4mo-9mo: *p*<0.0001; 4mo-12mo: *p*=0.0001; Tukey’s multiple comparisons) (Fig.1C). Additionally, we verified that DBA/2J IOPs were similar to age-matched controls at 4-months-old (*p=*0.96, Tukey’s multiple comparisons), and that DBA/2J IOPs were significantly elevated in comparison to controls at 9-and 12-months (9mo: 20.1±6.0 mmHg, *p*<0.0001; 12mo: 19.3±4.6, *p=*0.0001; Tukey’s multiple comparisons) (Fig.1C). As a measure of sustained IOP elevation, we calculated the area under the curve (AUC) of individual IOP series, which generated AUC(IOP) values for each eye, in mmHg*days. Using these measurements, we found that left eye and right eye IOP measurements were tightly correlated with each other for individual mice [Figure 1D; F(1,73)=2043, R^2^=0.97, p<0.0001]. These data verify that age-related OHT is conserved within our DBA/2J colony and demonstrate that our age groups are appropriate for comparisons of prolonged IOP exposure.

**Figure 1.**
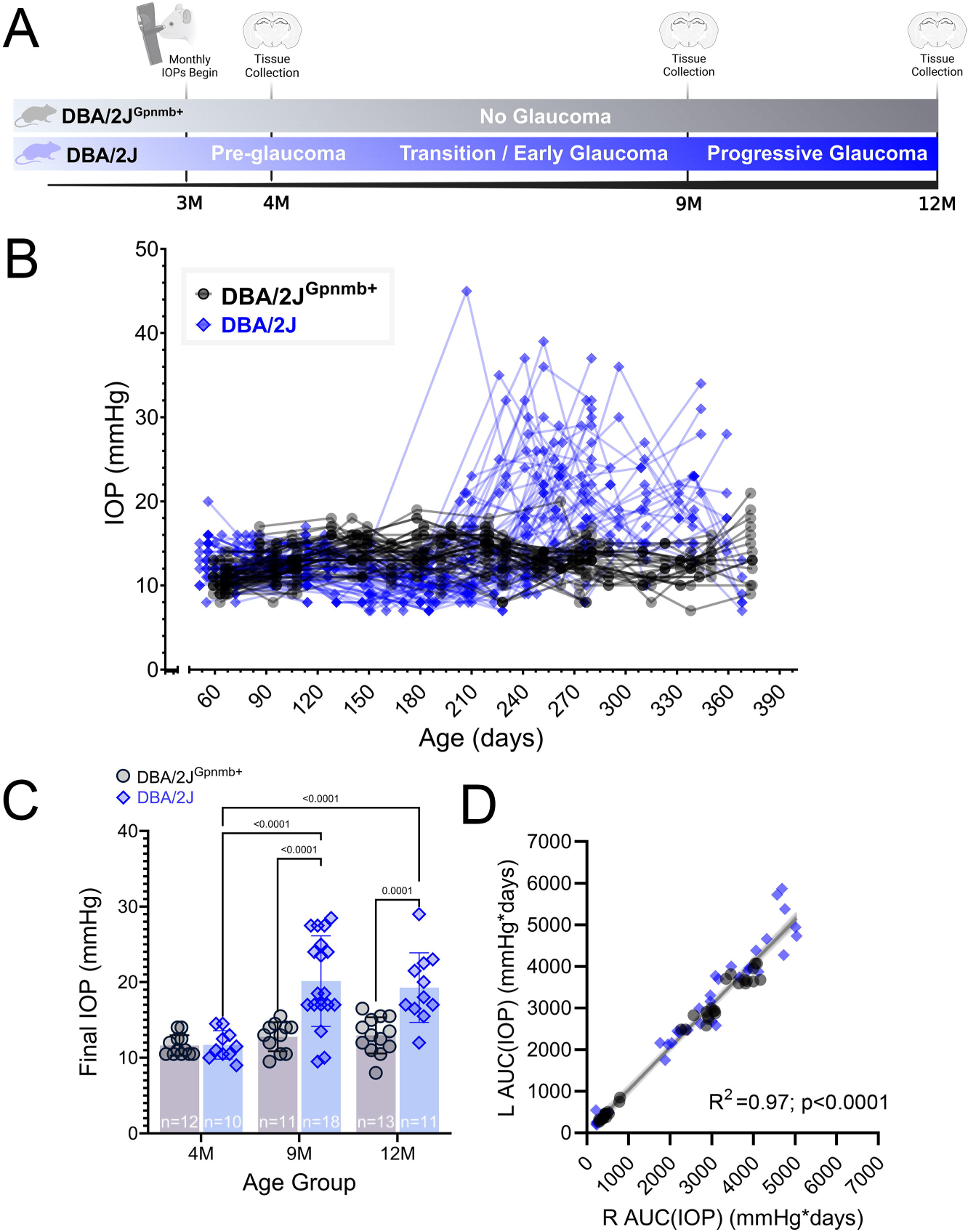
Glaucoma-relevant age groups are stratified in accordance with the variability and onset of DBA/2J ocular hypertension. **(A)** Experimental design indicating timing of intraocular pressure (IOP) measurements and tissue collection. **(B)** IOP measured in individual eyes (left and right) from mice included in the present study. **(C)** Group data of final left eye IOP measurements taken before tissue collection. The p-values represent pairwise Tukey’s multiple comparison tests following two-way ANOVA. **D)** Scatter plot and linear regression of integrated IOP [AUC(IOP)] from left and right eyes. Shaded region represents 95% confidence interval.

To determine the impact of OHT on RGC inputs to the dLGN throughout the progression of DBA/2J glaucoma, we stained coronal dLGN sections with an antibody against vesicular glutamate transporter 2 (vGluT2) and quantified vGluT2+ puncta within a central region of the dLGN core as a measure of RGC axon terminal density [17,19] (Fig. 2A). No age-related differences were detected between the mean vGluT2+ puncta densities of control samples (4mo-9mo: *p*=0.85; 9mo-12mo: *p*=0.94; Šídák’s multiple comparison test) (Fig. 2B). In contrast, when comparing DBA/2J vGluT2+ puncta densities across ages, we observed a significant reduction at the 12-month time point (4mo-12mo: *p*<0.0001; 9mo-12mo: *p*=0.020; Šídák’s multiple comparison test). Although the difference between 4-month and 9-month DBA/2J samples did not reach significance (*p*=0.066; Šídák’s multiple comparison test), both 9- and 12- month-old DBA/2J samples exhibited significantly lower vGluT2+ puncta densities than age-matched controls (9m: *p*=0.0006; 12m: *p*<0.0001; Šídák’s multiple comparison test) (Fig. 2B).

**Figure 2.**
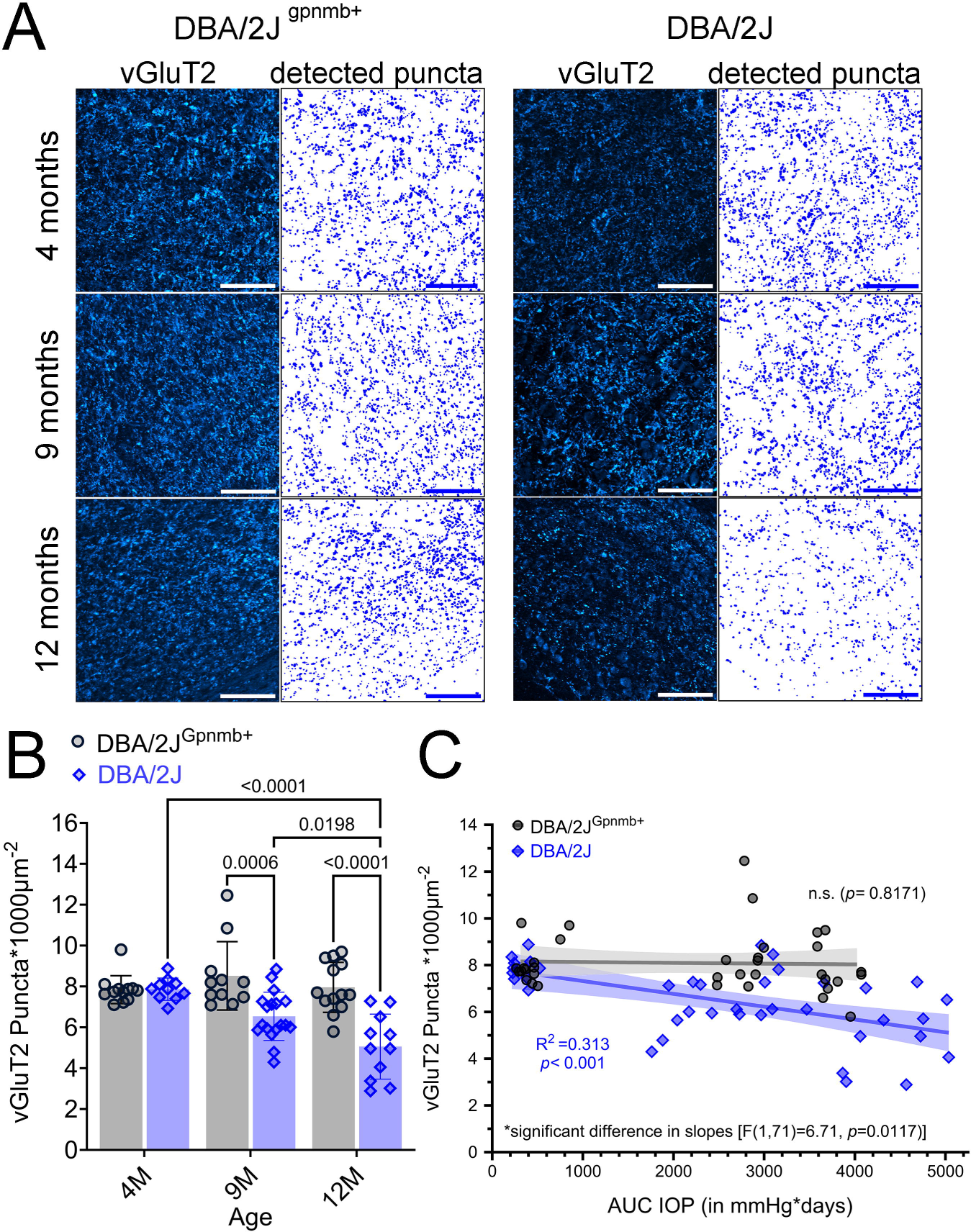
Age- and IOP-associated loss of vGluT2+ puncta in the DBA/2J dLGN. **(A)** 2- photon microscopy images of vGluT2+ RGC axon terminals from dLGN core and detected puncta. vGlut2+ dLGN TC neuron somas are more apparent with decline in punctate RGC axon terminal staining in 9- and 12-month-old DBA/2J images. Scale bar = 100 mm. **(B)** Group data of mean vGluT2+ densities factored by age and strain along with the results of significant (p<0.05) pairwise Šidák’s multiple comparison test comparisons. **(C)** Scatter plot of vGluT2+ puncta densities as a function of AUC(IOP) for each strain. Simple linear regressions were used to test associations between plotting factors and slopes from each genotype differed. Shaded area around regression line represents the 95% confidence interval.

Given the glaucoma-specific reduction in DBA/2J RGC-inputs, we next asked whether there was an association between OHT and vGluT2+ puncta density. We found that dLGN vGluT2+ density was negatively correlated with AUC(IOP) in DBA/2J mice (R^2^=0.313, *p*=0.0002). For DBA/2J^Gpnmb+^ control mice, in contrast, there was no significant correlation of vGluT2 density with AUC(IOP) (*p*=0.82). Strain-specificity was confirmed by slope comparison [*F*(1,71)=6.71, *p*=0.012; Fig. 2C].

To investigate whether vGluT2+ loss coincides with an increase in the presence of molecular machinery related to synaptic pruning, we stained a subset of DBA/2J (n=25) and DBA/2J^Gpnmb+^ (n=22) dLGN tissue sections with an antibody against complement component C1q (Fig. 3A), which is a molecular tag that facilitates the removal of excess retinogeniculate synapses during development [43,48,49]. Results from pairwise comparisons of mean C1q- peak-intensities revealed that C1q expression within the dLGN was low and unchanging for DBA/2J^Gpnmb+^ controls across all time points (4mo-9mo: *p*=0.79; 9mo-12mo: *p*>0.99; 4mo-12mo: *p*=0.52; two-tailed Šidák-corrected multiple comparisons tests). In contrast, there was an age-related increase in the mean of C1q-peak-intensities of DBA/2J dLGNs (4mo-9mo: *p*=0.025; 9mo-12mo: *p*=0.0034; 4mo-12mo: *p*<0.0001; two-tailed Šidák multiple comparisons tests) that could be statistically distinguished from age-matched controls by 12-months-old (*p*<0.0001, Šidák multiple comparisons test) (Fig. 3B). Given the timing of the C1q increase observed in DBA/2J samples, we hypothesized that C1q values might be associated with OHT, so we paired C1q-peak-intensities with their corresponding AUC(IOP) values and performed simple linear regressions (Fig. 3C). Indeed, C1q labeling intensity significantly correlated with AUC(IOP) in the dLGN of DBA/2J mice [*F*(1,30)=18.7, *p*=0.0002], but not in DBA/2J^Gpnmb+^ controls [*F*(1,23)=0.999, *p*=0.33]. Strain-specificity was upheld by results of slope comparison [*F*(1,53)=7.37, *p*=0.0089] (Fig. 3C). We next asked whether C1q-peak-intensities corresponded to the apparent loss of vGluT2+ RGC axon terminals in the dLGN, finding a negative correlation of vGluT2+ puncta density with C1q labeling intensity for DBA/2J mice [*F*(1,30)=13.1, R^2^=0.304, *p*=0.0011], whereas there was no such significant relationship for the DBA/2J^Gpnmb+^ controls (Fig. 3D). Despite the difference, no distinction could be made between the regressions between strains [*F*(1,53)=3.21, *p*=0.079] (Fig. 3D).

**Figure 3.**
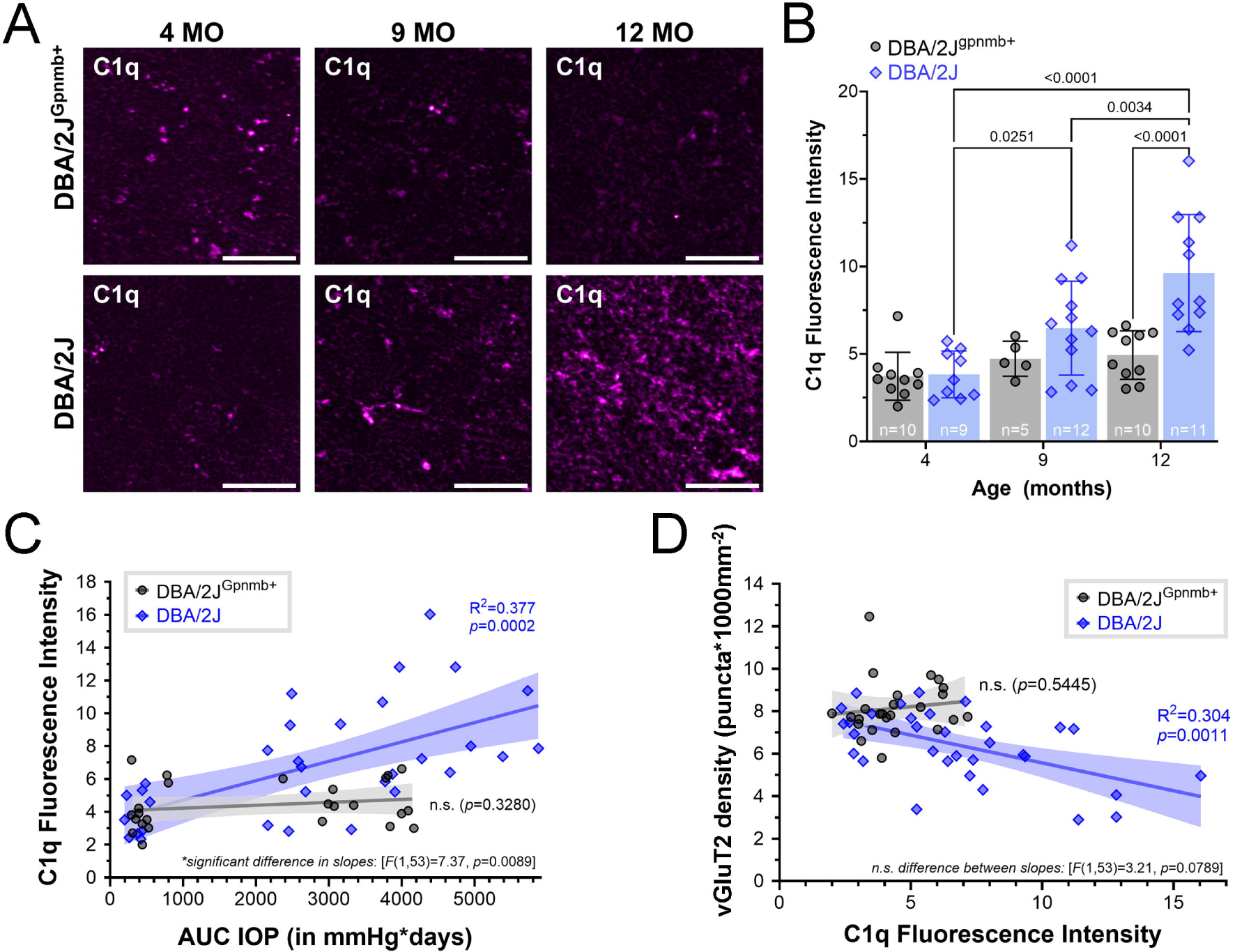
C1q intensity increases as a function of age and IOP in DBA/2J dLGNs. **(A)** Representative images of anti-c1q immunostaining of dLGN tissue obtained from 4-, 9-, and 12- month-old (MO) DBA/2J and DBA/2J^Gpnmb+^ mice (scale bar = 20 µm). **(B)** Bar graph factoring age and strain into peak C1q fluorescence intensities that were measured within the highest- expressing plane of C1q-stained dLGN volumetric images. Bar height and error bar represent the mean SD. Sample size (number of mice) for each condition are located within bars. Significant (p<0.05) pair-wise comparisons (two-tailed, Šidák-corrected multiple comparison tests) are shown with p-values. **(C)** Individual C1q intensities plotted against their corresponding IOP integrals for each strain, overlain with line fits and 95% confidence intervals resulting from simple linear regressions and follow-up slope comparisons. **(D)** Scatterplot of C1q intensity as a function of vGluT2+ puncta density in DBA/2J and DBA/2J^Gpnmb+^ dLGNs with linear regression outputs and follow-up slope comparison results.

We next sought to test whether IOP and/or age impacted the number or morphology of microglia in the dLGN, predicting that IOP-triggered pathology would evoke a shift of microglia morphology toward a more “active” state. Therefore, we obtained DBA/2J and DBA/2J^Gpnmb+^ dLGN tissue at three pathologically-relevant ages for the purpose of characterizing the microglial response to OHT in the dLGN throughout the progression of DBA/2J glaucoma. This was tested by performing a variety of morphometric analyses on Iba1+ cells within the dLGN to quantify features relevant to microglial ramification. A two-way ANOVA testing the effects of age and strain on the number of dLGN-present microglia resulted in significant main effects of age [*F*(2,69)=7.996, *p*=0.0008] and strain [*F*(1,69)=12.53, *p*=0.0007] on Iba1+ cell counts, without a significant interaction [*F*(2,69)=1.151, *p*=0.3324]. Regardless of strain, Iba1+ cell counts from 4- month-old samples showed that, on average, the central dLGN contained 23±4 microglia in our imaging volume, or 4200±730 microglia/mm^3^. Despite the lack of interaction, pairwise comparisons between Iba1+ cell counts in DBA/2J^Gpnmb+^ controls failed to uphold the main effect of age [4mo-9mo: *t*(69)=1.279, *p*=0.4980; 9mo-12mo: *t*(69)=0.7996, *p*=0.9872; 4mo-12mo: *t*(69)=2.152, *p*=0.1011], as the 24±4 average at 9-months only increased to 26±4 at 12-months (Fig.4B). In the DBA/2J dLGN, microglia presence was increased at both glaucoma-relevant ages (9mo: 31±7; 12mo: 30±6) in comparison to the 4-month-old pre-glaucoma condition (24±3): 4mo-9mo [*t*(69)=3.56, *p*=0.0021; 4mo-12mo, *t*(69)=2.95, *p*=0.0130 (Šidák multiple comparison; Fig.4B]. Although high microglia counts persisted within DBA/2J dLGNs at 12- months (30±6) compared to 4-months, *t*(69)=2.95, *p*=0.130, they did not differ from 9-month counts [*t*(69)=0.2986, *p*=0.9872 (Šidák multiple comparison; Fig.4B)].

**Figure 4.**
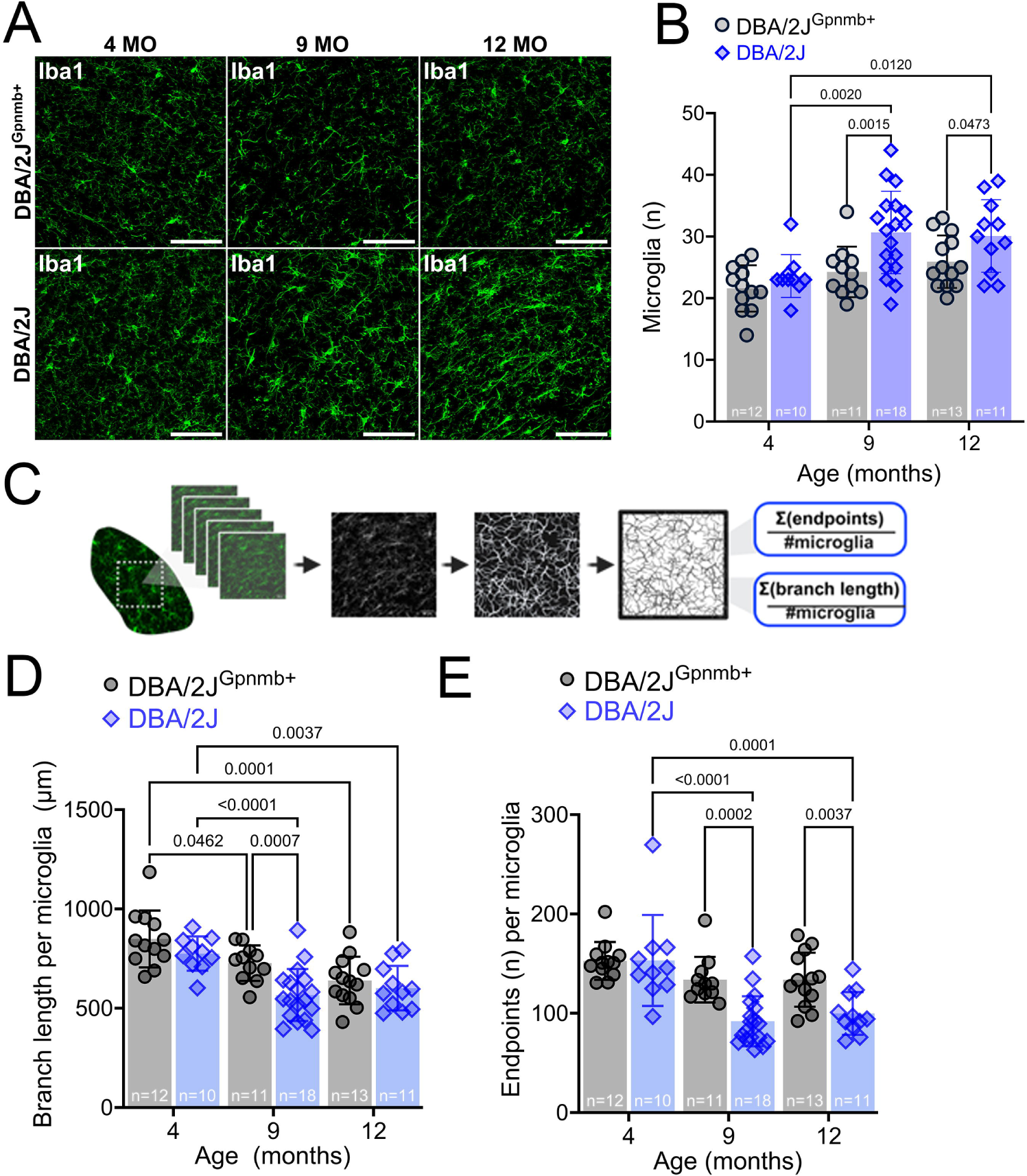
Morphology of DBA/2J dLGN-resident microglia is polarized at glaucoma- relevant time points. **(A)** Representative maximum intensity projections of 2-photon-acquired image stacks (z-depth: 40µm) for visualizing dLGN-resident microglia (Iba1 immunostained) in 4-, 9-, and 12-month-old DBA/2J and DBA/2J^Gpnmb+^ controls. Scale bar: = 100 µm. **(B)** Group data showing the number of microglia contained within each volumetric dLGN image stack for 39 DBA/2J (blue; diamonds) and 36 DBA/2J^Gpnmb+^ (gray, circles) mice. Two-way ANOVA testing resulted in significant main effects for age [*F*(2,69)=1.51, *p*=0.3224] and strain [*F*(1,69)=12.53, *p*=0.0007] on mean microglia counts, with a nonsignificant interaction term [*F*(2,69)=1.51, *p*=0.3224]. Pairwise comparisons are p-values from Šidák-corrected multiple comparison tests. **(C)** Illustrative workflow showing skeletonization of Iba1 images and analysis of total microglia branch lengths and quantification of endpoints to investigate microglia morphology. **(D)** Group data of microglia branch length from the skeleton analysis. Results from two-way ANOVA, revealed age effects [*F*(2,69)=17.6, *p*<0.0001] and strain effects [*F*(1,69)=10.7, *p*=0.0016] without significant interaction [*F*(2,69)=1.82, *p*=0.1695]. Pairwise comparisons are p-values from Šidák-corrected multiple comparison tests. **(E)** A two-way ANOVA generated a significant interaction term for the effects of age and strain on microglia endpoint averages [*F*(2,69)=3.889, *p*=0.0251] along with significant main effects: age [*F*(2,69)=14.88, *p*<0.0001], strain [*F*(1,69)=15.05, *p*=0.0002]. Pairwise comparisons are p-values from Šidák-corrected multiple comparison tests.

Next, we assessed microglial ramification by quantifying microglia cell branch length and endpoint averages from global analysis of skeletonized Iba1 fluorescence images [45] and performing similar analyses with respect to age and strain (Fig.4C-F). Descriptions of arbor complexity are a powerful tool for characterizing the state of microglia polarization within a neurodegenerative context because microglia exist along a morphological continuum that ranges from intricately branched under homeostatic/surveilling conditions to ameboid such as during immune activation.

In DBA/2J^Gpnmb+^ and DBA/2J dLGN tissue sections, there appeared to be both age- and genotype-dependent effects on branch length per microglia cell based on a 2-way ANOVA results indicating that 28.2% of Iba1+ branch length variance could be explained by age [*F*(2,69)=17.6, *p*<0.0001], whereas only 8.6% could be explained by genotype, (*F*(1,69)=10.7, *p*=0.0016), without significant interaction, [*F*(2,69)=1.82, *p*=0.1695]. Šidák multiple comparisons tests further revealed reduced branch length at 9 and 12 months in both DBA/2J and DBA/2J^Gpnmb+^ samples. Divergence of branch length was also apparent in comparisons of 9- month-old DBA/2J^Gpnmb+^ and DBA/2J, with a lower branch length apparent in DBA/2J (p=0.0007), pointing toward an acceleration of the age-dependent departure from the ramified, homeostatic phenotype in DBA/2J mice (Fig.4D).

Analysis of the number of microglia endpoints per cell, another measure of microglial complexity, revealed a similar pattern (Figure 4E), with a 2-way ANOVA revealing significant effects of both age [*F*(2,69)=14.88, *p*<0.0001] and genotype [*F*(1,69)=15.05, *p*=0.0002]. Unlike for branch length, analysis of endpoints revealed a significant interaction term for the effects of age and strain on microglial endpoint averages [*F*(2,69)=3.889, *p*=0.0251]. Although Šidák multiple comparison tests revealed no significant difference between 4-, 9- and 12-month old DBA/2J^Gpnmb+^ microglia, there was a lower number of microglia process endpoints per cell in the 9- and 12-month old samples compared to 4-month. Additionally, comparisons of DBA/2J- gpnmb+ with DBA/2J at 9-month ages revealed lower numbers of microglia endpoints, again indicating a simplified morphology consistent with a move away from the surveilling homeostatic phenotype.

We next sought to determine whether OHT and loss of RGC axon terminals might be related to microglial ramification within the visual thalamus, since these cells rapidly respond to changes within synaptic microenvironments. We tested this by performing linear regressions to assess potential relationships of OHT, RGC axon terminal density, and complement C1q with microglia morphology within the visual thalamus (Fig.5A-C). We observed negative associations between AUC(IOP) and measurements of Iba1+ cell complexity (Fig. 5A); for both DBA/2J^Gpnmb+^ controls and DBA/2J mice, Iba1+ cell branch length was negatively correlated with AUC(IOP) (Figure 5A). Likewise, the number of microglia endpoints was negatively correlated with IOP for both DBA/2J^Gpnmb+^controls and DBA/2J Iba1+ cells. For both endpoints and branching, slope comparison results showed that IOP-associations were strain-independent [branching: *F*(1,71)=2.19, *p*=0.1429; endpoints: *F*(1,71)=0.65, *p*=0.4229].

**Figure 5.**
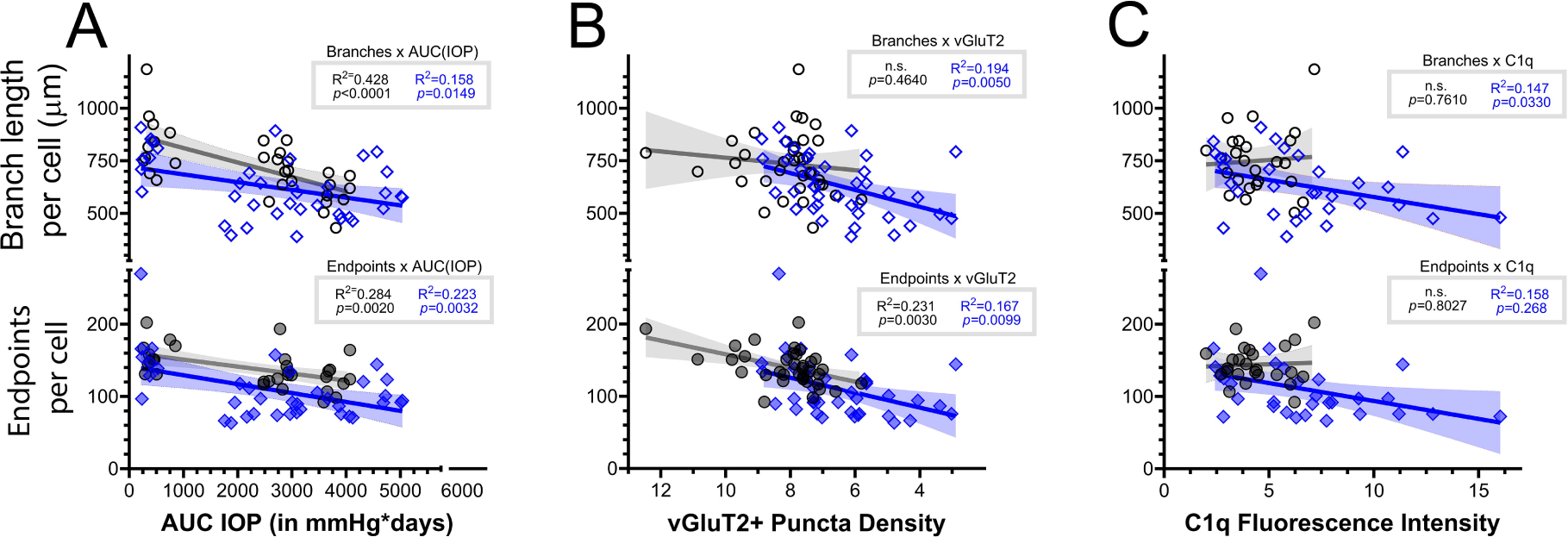
DBA/2J glaucoma is accompanied by age- and OHT-associated polarization of Iba1+ cells within the visual thalamus. **(A)** Scatterplots and linear regressions of microglia branch length and endpoints per cell from skeleton analysis plotted against AUC(IOP). Shaded area represents the 95% confidence interval. Black circles: DBA/2J^Gpnmb+^ controls. Blue diamonds: DBA/2J. **(B)** Scatterplots and linear regressions, as in **A** of microglia skeleton parameters plotted against vGluT2 puncta density. **(C)** Scatterplots and linear regressions of microglia parameters plotted against C1q fluorescence intensity.

We next assessed whether features of Iba1+ complexity were linearly associated with vGluT2+ puncta density. Indeed, we found a negative association between Iba1+ branch length and vGluT2+ puncta for DBA/2J mice, but not for DBA/2J^Gpnmb+^ controls. We confirmed the relationship of AUC(IOP) and vGlut2 puncta density to be independent of genotype by comparing the slope outputs of strain-specific regressions, finding no significant difference in slopes from each regression [F(1,71)=1.15, p=0.29]. Regarding microglial endpoints per cell, we found that branch length was negatively associated with vGluT2+ puncta densities in both DBA/2J mice and DBA/2J^Gpnmb+^ controls (Fig.5B). Additionally, there was no significant difference in slopes from each regression [F(1,71)=0.03, p=0.86]. We found that C1q intensities were useful for predicting Iba1+ cell complexity in the dLGNs DBA/2J but not in DBA/2J^Gpnmb+^ controls; DBA/2J Iba1+ cell branching and endpoint measurements both decreased as a function of C1q intensity. There was no such relationship for DBA/2J^Gpnmb+^ controls. Despite this, we did not detect a significant difference in the slopes of each regression (F(1,52)=1.24, p=0.27). We found a similar pattern for measurements of Iba1+ cell branch length, with a significant relationship of branch length and C1q intensity for DBA/2J, but not DBA/2J^Gpnmb+^ controls. Additionally, we did not detect a significant difference in slopes of each regression (F(1,52)=1.07, p=0.30).

In addition to taking on simplified/ameboid morphologies, microglia can respond to central nervous system injury by taking on an elongated/rod/bipolar-like morphology [39–42]. These rod microglia might play important roles in phagocytosis of cellular debris and/or repair of CNS circuits. Qualitatively, we observed that many Iba1-labeled cells were elongated and aligned in a common direction in dLGN sections from aged DBA/2J mice, suggesting that OHT might trigger adoption of a rod-like morphology in the dLGN (Fig 4A). To quantitatively analyze microglia morphology and test this, performed a fractal analysis of individual microglia cells. We randomly selected four individual microglia cells from each of our multiphoton image stacks and reconstructed 2-dimensional binarized projections of their somas (Figure 6A). Skeleton analysis of these individual microglia generally affirmed results from the global analysis (Figure 6 B&C), showing a lower number of endpoints and lower total branch length in the 9 month-old DBA/2J compared to 4 month-old DBA/2J, although we did not detect significant differences when compared to age matched controls, which likely results from variability and inclusion of only four individual microglia cells per mouse. We next used the FracLac ImageJ plug-in to analyze cell perimeter, circularity, fractal dimension (*D_f_*), and span ratio of each cell (Figure 7). We hypothesized that microglia perimeters would decrease with simplifying of their morphologies with age and in ocular hypertensive dLGN, but perimeter was indistinguishable between all conditions (Fig. 7A), as revealed by a nested one-way ANOVA (*F(*5,48)=1.013, *p*=0.4025).

**Figure 6.**
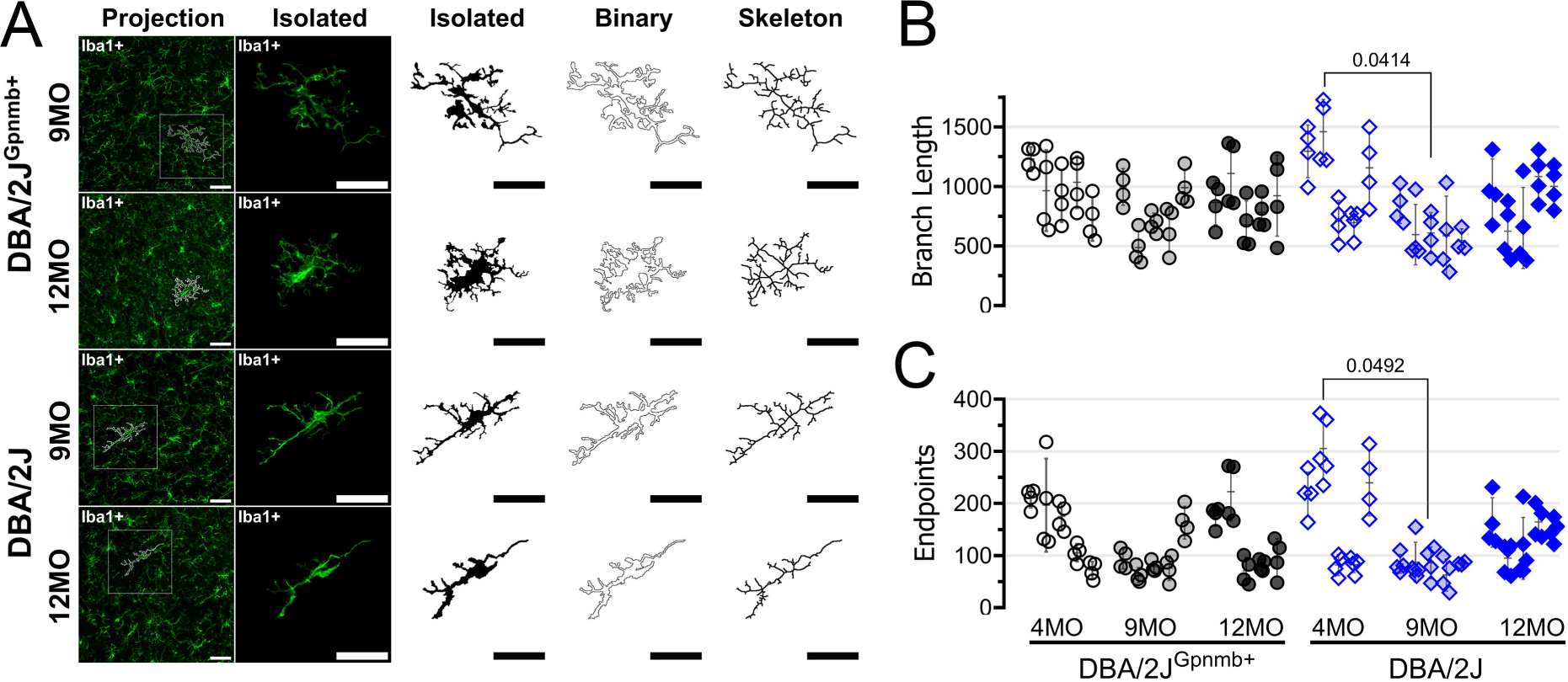
Analysis of individual microglia morphology. **(A)** Example images and reconstruction of randomly-selected Iba1 immunolabeled microglia from 9- and 12-month old (MO) dLGN sections from DBA/2J^Gpnmb+^ and DBA/2J mice. **(B&C)** Skeleton analysis of branch length and endpoints of four individual microglia cells per mouse. Each column of data points contains four cells from a single animal. Pairwise comparisons represent significant (p<0.05) p- values from Šidák-corrected multiple comparison tests.

**Figure 7.**
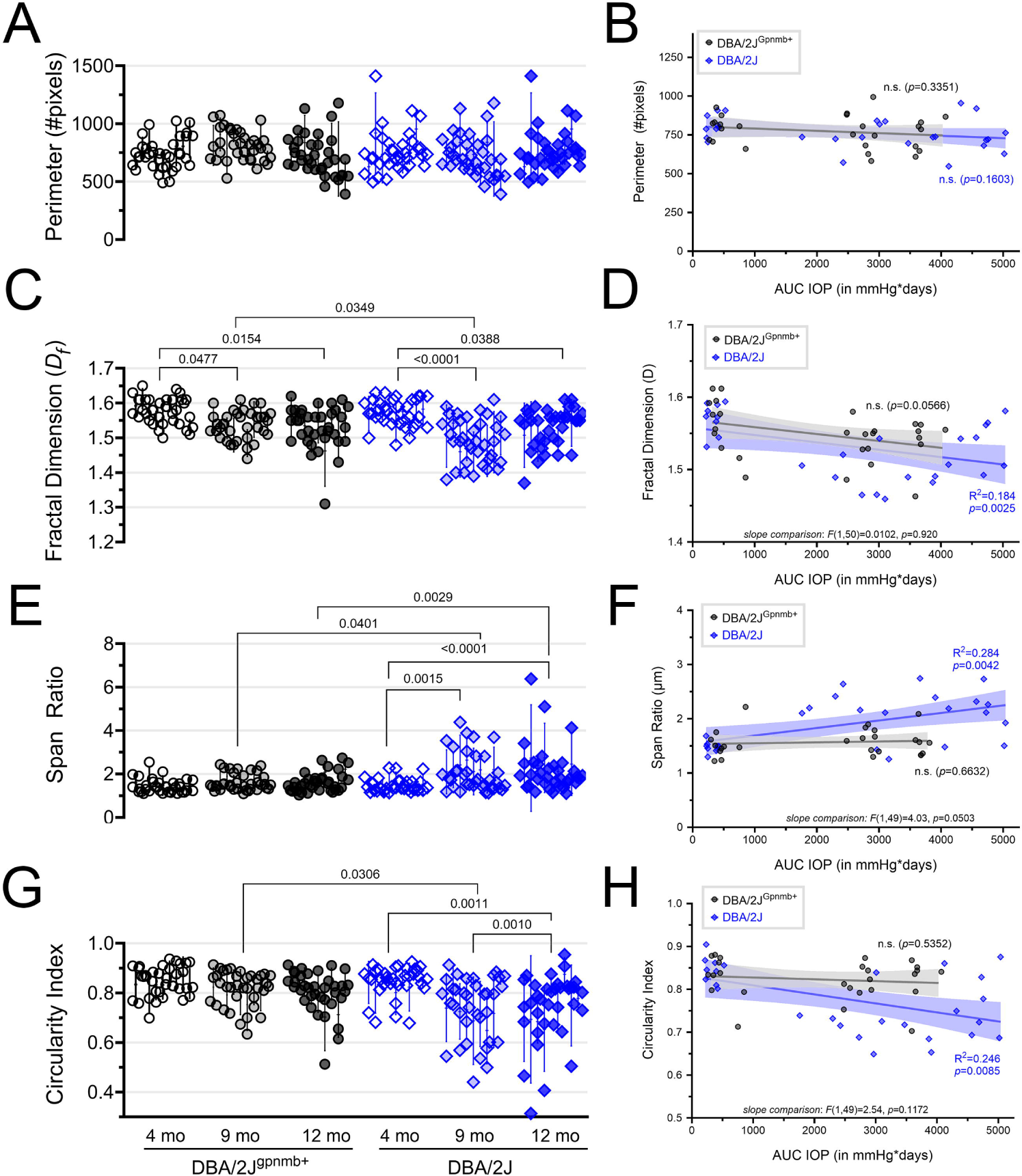
Fractal analysis of microglia morphology reveals a shift toward rod-like microglia morphology in aged DBA/2J mice. **(A)** Plot of microglia perimeter (# pixels) output of the FracLac analysis from four randomly selected and manually-reconstructed microglia cells per mouse of DBA/2J^Gpnmb+^ controls (black circles) and DBA/2J mice (blue diamonds) at 4, 9, and 12 months of age. Each column of data points corresponds to measurements from a single animal. **(B)** Scatterplot and simple linear regression of microglia perimeter measurements averaged from individual animals plotted against AUC(IOP). Shaded area represents the 95% confidence intervals. **(C&D)** Similar to **A** and **B**, plotting the FracLac parameter “Fractal Dimension” (*D_f_*). Pairwise comparisons represent significant (p<0.05) results of Šidák multiple comparison tests following a nested one-way ANOVA (p<0.0001). **(E-H)** Similar analyses as A-D for FracLac parameters “Span Ratio” and “Circularity Index”.

Accordingly, we found that perimeter of individual microglia cells did not significantly correlate with AUC(IOP) for either strain (DBA2J: *F*(1,24)=0.968, *p*=0.335; DBA/2J^Gpnmb+^ controls: *F*(1,25)=2.09, *p*=0.1603) (Fig.7B).

Fractal dimension (*D_f_*) provides a general measure of microglia process complexity and this value revealed a pattern that mirrored measurements we obtained using skeleton analyses of process length and endpoints (Fig. 4). Specifically, we found a reduction in *D_f_* with increasing age in both DBA/2J^Gpmnb+^ controls and DBA/2J mice (one-way ANOVA), revealing significant differences between 9-month control and DBA/2J measurements. Likewise, *D_f_* was negatively correlated with AUC(IOP) for DBA/2J mice. A similar regression analysis for the DBA/2J^Gpnmb+^ control population had a p-value of 0.057 and comparison of the slopes between genotypes showed they were similar (F(1,50)=0.01, p=0.92). This result, which examines microglia complexity without consideration of shape, aligns closely with our findings from the global skeleton analysis, highlighting an apparent age- and IOP-dependent reduction in microglia in both DBA/2J and DBA/2J^Gpnmb+^ controls with more pronounced effects observed in DBA/2J at the 9-month time point.

To specifically examine microglia elongation and test whether they adopt a more bipolar/rod morphology in the dLGN in response to OHT, we analyzed two parameters: 1) the span ratio, which is the ratio of the longest axis to the orthogonal axis of a convex hull drawn around the microglial cell, with higher values indicating a more elongated morphology, and 2) circularity index, which quantifies how close the convex hull is to a circle on a scale of 0-1, with 1 being circular. For span ratio, microglia cells from older DBA/2J tissue sections had higher values and a one-way nested ANOVA revealed a significant difference among groups. Pairwise analyses using Šídák’s multiple comparisons tests showed that DBA/2J microglia span ratios were significantly higher than controls at both glaucoma-relevant time points (9mo: p=0.0401; 12mo: p=0.0028) and increased with age in the DBA/2J population while span ratios were unchanged across age for the DBA/2J^Gpnmb+^ controls. Span ratio also significantly correlated with AUC(IOP) in the DBA/2J population, but not the DBA/2J^Gpnmb+^ controls, although comparisons of the slopes of the two regressions had a p-value of 0.050. Analysis of circularity index data revealed a similar pattern; following a significant nested one-way, ANOVA F(5,48)=7.514, p<0.0001, results of two-tailed Šídák-corrected t-tests showed lower circularity index values of cells from older DBA/2J tissue sections, revealing a loss of circularity compared to age-matched controls (p=0.0306). An age-related loss of microglial circularity was detected by comparisons of DBA/2Js to their 9- (p=0.0011) and 12-month-old (p=0.0010) counterparts. In a similar fashion to span ratio, the circularity index negatively correlated with AUC(IOP) in the DBA/2J population, but not the DBA/2J^Gpnmb+^ controls, although comparison of the slopes of the two regressions had a p-value of 0.12. This analysis highlights an adoption of elongated/rod-like morphology of dLGN microglia with increasing cumulative IOP specifically in DBA/2J mice and not DBA/2J^Gpnmb+^ controls.

To further test for impacts of elevated IOP, we performed bulk RNA-sequencing of microdissected dLGN tissue from 9 month-old DBA/2J and DBA/2J^Gpnmb+^ control mice. After monthly IOP measurements (Figure 8A), dLGN tissue from both right and left hemispheres from three animals per genotype was pooled to give five DBA/2J samples (total = 15 mice) and three DBA/2J^Gpnmb+^ samples (total = 9 mice). We identified a total of 77 upregulated and 7 downregulated differentially expressed genes (DEGs; Figure 8B) that met criteria for differential expression (≥1.5 fold-change, q<0.05). Roles of these genes were identified using a gene ontology (GO) analysis and Kyoto Encyclopedia of Genes and Genomes (KEGG) enrichment analysis tools. Among the pathways identified in GO analysis were several related to immune system function, including “immune response” (GO:0006955), “innate immune response” (GO:0045087), “antigen processing and presentation” (GO:0019882), etc. Likewise, top results on KEGG analysis included “antigen processing and presentation”, “phagosome”, and “natural killer cell mediated cytotoxicity”. Upregulated protein-coding genes included major histocompatibility complex (MHC) class I and class II genes (*H2-Q2*, *Cd74,* and *H2-Ea*), and CD11c*/Itgax* of the complement pathway, consistent with upregulated immune system function in DBA/2J dLGN. CD11c/*Itgax* expression is characteristic of a subpopulation of microglia and supports phagocytosis of cellular and synaptic debris. Galectin-3 (*Lgals3*) was upregulated 4.8- fold and plays roles in microglia activation and inflammatory responses. Expression of two neurotrophins, neurotrophin-3 and neurotrophin-5 (*Ntf3* and *Ntf5*) was also significantly upregulated. These can be expressed by microglia and regulate neuron survival as well as microglia activation and proliferation and interact with MHCII pathways.

**Figure 8.**
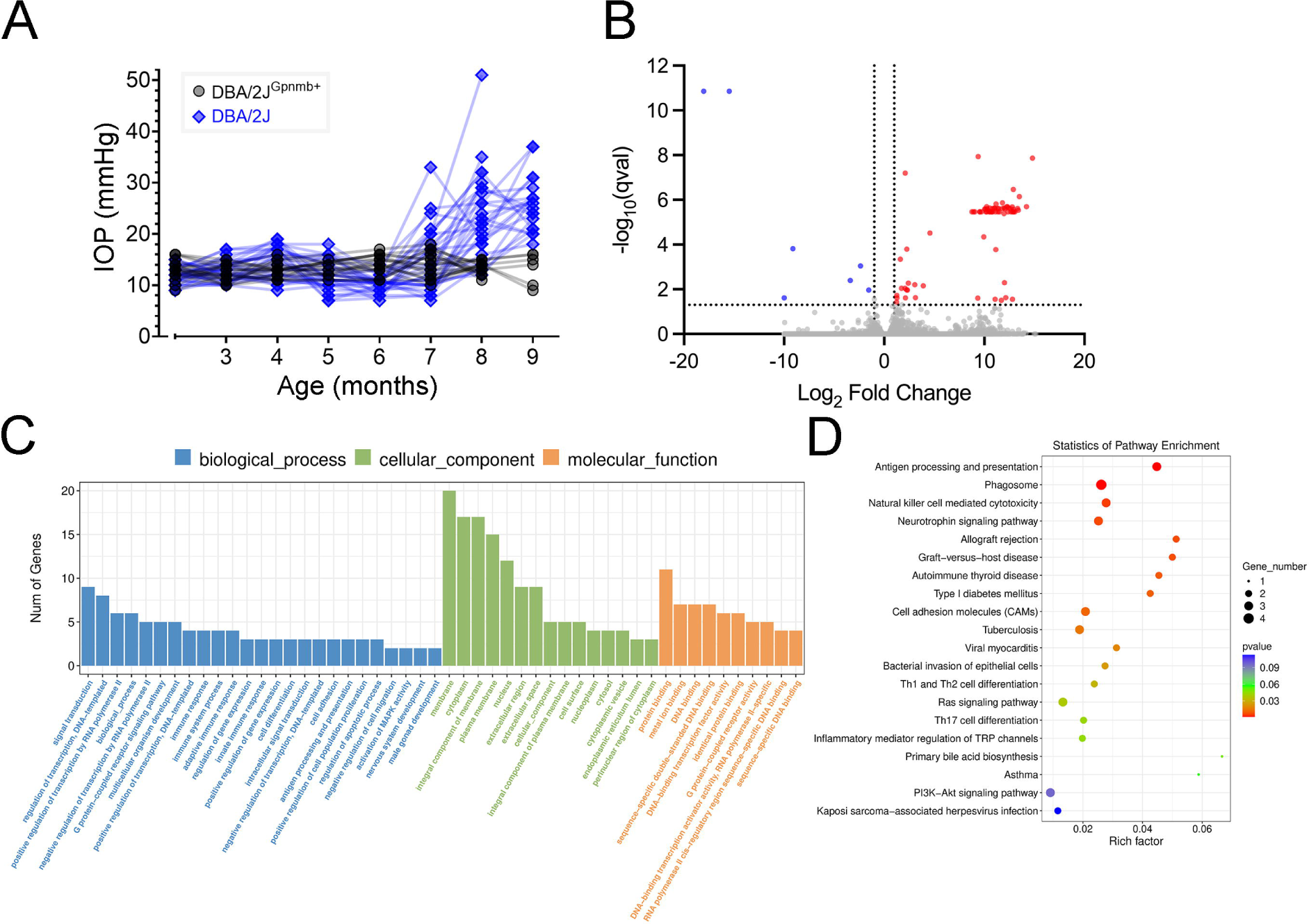
Alterations in gene expression in the dLGN of DBA/2J mice revealed by RNA sequencing. **(A)** Intraocular pressure measurements from DBA/2J^Gpnmb+^ (n = 18 eyes, 9 mice) and DBA/2J (n = 30 eyes,15 mice. **B**) Volcano plot of 33,064 genes showing that 77 were significantly upregulated and 7 that were significantly downregulated in the DBA/2J dLGN. Significantly up- or down-regulated genes were identified by criteria of a 1.5-fold change and false discovery rate-corrected p value < 0.05. **C)** Gene ontology enrichment analysis of differentially expressed genes. **D)** Kyoto Encyclopedia of Genes and Genomes (KEGG) enrichment analysis.

## Discussion

The goal of this study was to determine whether eye pressure and glaucoma influence microglia morphology in the dLGN, which is a critical RGC projection target for conscious vision. Use of anti-Iba1 immunofluorescence and analysis of microglia numbers and morphology point to an IOP-related increase in microglia number in the dLGN. These microglia appeared to "activate”, taking on a less ramified, simpler morphological phenotype with increasing age in both mice with elevated IOP and in non-ocular hypertensive controls [47]. However, this age-related de-ramification was accelerated in the dLGN of DBA/2J mice and correlated with C1q labeling and vGluT2+ RGC terminal loss, pointing to a confluence of age and glaucomatous pathology impacting dLGN microglia. In addition to the apparently earlier onset of microglia de- ramification, analysis of individually-reconstructed microglia using a fractal analysis showed that microglia in the DBA/2J dLGN adopted a “rod-like” elongated [39–42,46] morphology that was related to IOP and was not apparent in the control population. Finally, transcriptomics analysis of dLGN tissue from 9-month-old DBA/2J mice revealed upregulation of numerous genes related to immune signaling and implicated in microglia function.

Morphology is typically viewed as a marker of microglial functional state with complex/ramified microglia actively monitoring their local environment and responding to stressors by progressing toward a simplified/de-ramified morphology [50,51]. Our results from a global skeleton analysis of microglia branch length and endpoints point to an effect of age triggering a shift of microglia morphologies toward this more de-ramified morphology. In mice with glaucoma, however, this process was accelerated, and this was related to cumulative IOP measurements. This suggests that age alone might contribute to some level of microglia de- ramification in the dLGN in the absence of obvious visual system pathology, but that this process is accelerated or more pronounced in the disease context. We also found a relationship of vGluT2-labeled RGC axon terminal loss and microglia morphology. This relationship was similar in DBA/2J and controls when we examined microglia endpoints but showed a distinction between DBA/2J and controls in analyses of microglia branch length. This implies that presynaptic integrity in the dLGN could be compromised by loss of microglial interaction, even in the absence of underlying disease, and highlights a potential threat of continuous microglial responses to IOP elevation. There was also a relationship of both measures of microglia morphology with complement cascade C1q expression only in DBA/2J dLGN and not control tissue. C1q is a key member of the classical complement cascade [43] and is involved in tagging CNS synapses for elimination, including in the developing dLGN [43,49]. Alternatively, microglia might not be the drivers of synapse elimination in the glaucomatous dLGN, but might instead be responding to RGC axon degeneration and clearing degenerated debris [48,49,51].

Using a fractal analysis of microglia morphology [45–47], we found that chronic IOP elevation caused numerous microglia in the dLGN to assume rod-like phenotypes, defined by thin, elongated somas with few protrusions [39,46,52,53]. The rod-like microglia morphology we report here resembles that seen in the somatosensory cortex barrel field following a diffuse traumatic brain injury [46,52,54]. Unlike the general de-ramification seen in aged dLGN and, at younger age in ocular hypertensive dLGN, rod morphology appeared to be unique to the ocular hypertensive DBA/2J dLGN; the slopes of linear regressions comparing two measures of microglia elongation – span ratio and circularity index – showed clear divergence between DBA/2J and DBA/2J*^Gpnmb+^*(Figure 7) whereas this was not the case for analyses that merely probed microglia process complexity without regard for cell shape (endpoints & branch length in Fig 5; *D_f_*in Fig 7). Prior work using rats with experimentally-elevated IOP showed apparent microglia de-ramification in dLGN and superior colliculus, although it is unclear if there were rod-like dLGN microglia in that study [55]. Optic nerve transection from enucleation also leads to dramatic microglia de-ramification, but without obvious rod microglia [56], suggesting that the IOP-triggered optic nerve injury in the DBA/2J dLGN favors microglial elongation. Within the DBA/2J dLGN, most rod microglia appeared oriented parallel to the optic tract, possibly implying that they align along damaged and/or degenerating axons coursing across the dLGN. Rod microglia are generally observed to align with neuron dendrites and axons [39,42,52] and several studies have already shown that optic nerve damage causes rod microglia to accumulate exclusively around RGC compartments in the retina [32–34,57]. Severe optic nerve injury resulted in trains of rod microglia that fanned away from the optic disc along the axons of injured RGC [58].

The unique functions of rod microglia are still debated and it is possible that rod microglia are not a single “type” and that a rod-like morphology is shared by numerous microglia classes with distinct functional roles. The elongated morphology and association with neuronal processes suggest they might provide structural support to stressed or damaged axons and/or dendrites. Restriction of activated microglia to sites of direct injury is a common finding and might imply a neuroprotective role [57,59]. Rod-like microglia might also be an intermediate morphological class, representing a step between ramified and “active” ameboid morphologies [40,51]. One cell culture study showed that rod-like microglia have reduced expression of both pro- and anti-inflammatory markers early in culture, but could be rapidly shifted to a pro- inflammatory/ameboid phenotype following exposure to lipopolysaccharides [53]. Rod microglia have also been hypothesized to be involved in synaptic stripping, whereby synapses are removed from damaged neurons, possibly in an attempt at preserving neurons by reducing metabolic stress [39,40,42]. Indeed, synaptic stripping might be occurring in the glaucomatous dLGN, as IOP-dependent loss of vGlut2-labeled RGC axon terminals is followed at later time points by a loss of TC neuron dendrites.

Overall, our transcriptomics data support a general picture of immune system activation in the dLGN of DBA/2J mice. GO and KEGG analyses of significantly up/down-regulated genes both revealed terms related to immune system function, antigen presentation, phagocytosis, etc. While we performed RNAseq on whole dLGN tissue samples, several upregulated genes have established ties to microglial function. For instance, *Cd74* was upregulated by approx. 4.3-fold while *H2-Ea* was upregulated by approx 4.6-fold. The H2-Ea protein is a major histocompatability complex II (MHC II) component while Cd74 (cluster of differentiation 74 protein) functions to regulate intracellular MHC II trafficking and membrane organization [60]. MHCII expression allows for surface presentation of antigens for recruitment of T-cells that can impact inflammatory processes and neurodegeneration [61]. MHC II expression by microglia is relatively low under basal conditions, but increases in microglia in response to disease states, meaning its expression can serve as a marker of microglia activation [61–63]. *Itgax*, which was upregulated approx. 26-fold, encodes the complement protein, Integrin alpha-X/CD11c, that is found on a neuroprotective population of microglia [64]. CD11c+ microglia are neuroprotective in development [64] and promote a regional bias towards myelin preservation [65,66]. In the developing brain CD11c+ is associated with a seven-fold increase in galectin-3 (*Lgals3*) expression [64]. We found a 4.8-fold increase in the Gal-3 gene *Lgals3* in the DBA/2J dLGN. Gal-3, which is a member of beta-galactoside binding lectin family [67,68], was also found to be upregulated in retinal microglia in an inducible rodent glaucoma study [55]. Gal-3 can be membrane-bound or secreted and its function is highly context-dependent. Gal-3 that is secreted by microglia after CNS injury suppresses proinflammatory cytokines and facilitates the removal of myelin debris, which in turn promotes effective remyelination [69]. In the mature brain, Gal-3-signaling occurs between microglia and injured oligodendrocytes during the remedial response to cuprizone-induced demyelination [70].

We also found a 3000-4000-fold increase in expression of Ntf3 and Ntf5, which encode Neurotrophin-3 (NT-3) and Neurotrophin-5 (NT-5). Neurotrophin expression can be upregulated in activated microglia [71,72] and serve a neuroprotective effect through activation of TrkC and TrkB receptors [73]. NT-3 signaling between microglia regulates proliferation and phagocytosis [71] and can influence MHC II expression [74]. Another neurotrophin, BDNF, can act through neuronal TrkB receptors, which are also activated by NT-3 and NT-5, to influence synaptic plasticity and neuronal excitability in the dLGN [75]. Although *Bdnf* expression was not altered in the DBA/2J dLGN, it is possible that microglial NT-3 and/or NT-5 mediate TC neuron homeostatic responses to IOP [17,19,76–79].

### Weaknesses of the current study

The current study focused on IOP-related alterations to microglia morphology in the dLGN. While morphology is clearly related to microglia function, we do not have a clear picture of what role microglia are playing in the glaucomatous dLGN. Future experiments depleting microglia from mice with glaucoma could help determine whether they ultimately play a supportive/neuroprotective role or whether microglia activation is a net detriment for dLGN responses to glaucoma. Such experiments are bound to be complicated, as a recent preprint showed disassembly of synapses in the retina following experimental IOP elevation was reduced by a microglial inhibitor [80], but 10-week-long systemic depletion of microglia in DBA/2J mice led to more severe optic nerve damage [31], suggesting that microglia might have divergent effects depending on disease stage or RGC compartment. Although we did not investigate whether rod-like microglia are dLGN-resident microglia that have transitioned morphologies or if they’re a population that has infiltrated, the stability of microglia numbers from 9- to 12-months suggests that they are resident microglia, as recruited macrophages are usually transient. Elsewhere, increases in rod microglia were reported to occur via local proliferation [53,58] rather than peripheral recruitment, but this would need to be tested under these conditions. Additionally, our RNA-sequencing relied on analysis of whole tissue isolates and pooling of tissue from several mice. This can introduce noise into the gene expression data, possibly hiding altered gene expression in specific dLGN cell types and masking the biological variability arising from the diverse IOP profiles in individual mice. Future work using either single-cell sequencing [29] or sequencing of microglia-enriched samples could help address this and allow for a clearer picture of microglia-specific IOP-related expression changes and functional implications.

## References

1. Flaxman SR, Bourne RRA, Resnikoff S, Ackland P, Braithwaite T, Cicinelli MV, et al. Global causes of blindness and distance vision impairment 1990-2020: a systematic review and meta-analysis. Lancet Glob Health. 2017;5: e1221–e1234. doi:10.1016/S2214-109X(17)30393-5

2. Lommatzsch C, van Oterendorp C. Current Status and Future Perspectives of Optic Nerve Imaging in Glaucoma. J Clin Med. 2024;13: 1966. doi:10.3390/jcm13071966

3. Quigley HA, Broman AT. The number of people with glaucoma worldwide in 2010 and 2020. Br J Ophthalmol. 2006;90: 262–267. doi:10.1136/bjo.2005.081224

4. Tham Y-C, Li X, Wong TY, Quigley HA, Aung T, Cheng C-Y. Global prevalence of glaucoma and projections of glaucoma burden through 2040: a systematic review and meta-analysis. Ophthalmology. 2014;121: 2081–2090. doi:10.1016/j.ophtha.2014.05.013

5. Weinreb RN, Aung T, Medeiros FA. The pathophysiology and treatment of glaucoma: a review. JAMA. 2014;311: 1901–1911. doi:10.1001/jama.2014.3192

6. Calkins DJ. Critical pathogenic events underlying progression of neurodegeneration in glaucoma. Prog Retin Eye Res. 2012;31: 702–719. doi:10.1016/j.preteyeres.2012.07.001

7. Calkins DJ. Adaptive responses to neurodegenerative stress in glaucoma. Prog Retin Eye Res. 2021;84: 100953. doi:10.1016/j.preteyeres.2021.100953

8. Seabrook TA, El-Danaf RN, Krahe TE, Fox MA, Guido W. Retinal input regulates the timing of corticogeniculate innervation. J Neurosci. 2013;33: 10085–10097. doi:10.1523/JNEUROSCI.5271-12.2013

9. Morrison JC, Johnson EC, Cepurna W, Jia L. Understanding mechanisms of pressure-induced optic nerve damage. Prog Retin Eye Res. 2005;24: 217–240. doi:10.1016/j.preteyeres.2004.08.003

10. Howell GR, Libby RT, Jakobs TC, Smith RS, Phalan FC, Barter JW, et al. Axons of retinal ganglion cells are insulted in the optic nerve early in DBA/2J glaucoma. J Cell Biol. 2007;179: 1523–1537. doi:10.1083/jcb.200706181

11. Nuzzi R, Dallorto L, Rolle T. Changes of Visual Pathway and Brain Connectivity in Glaucoma: A Systematic Review. Front Neurosci. 2018;12: 363. doi:10.3389/fnins.2018.00363

12. Yucel YH, Gupta N. A framework to explore the visual brain in glaucoma with lessons from models and man. Exp Eye Res. 2015;141: 171–178. doi:10.1016/j.exer.2015.07.004

13. Gupta N, Yücel YH. Brain changes in glaucoma. Eur J Ophthalmol. 2003;13 Suppl 3: S32–35.

14. Gupta N, Yücel YH. Glaucoma and the brain. J Glaucoma. 2001;10: S28-29. doi:10.1097/00061198-200110001-00011

15. Gupta N, Greenberg G, de Tilly LN, Gray B, Polemidiotis M, Yücel YH. Atrophy of the lateral geniculate nucleus in human glaucoma detected by magnetic resonance imaging. Br J Ophthalmol. 2009;93: 56–60. doi:10.1136/bjo.2008.138172

16. Gupta N, Ang L-C, Noël de Tilly L, Bidaisee L, Yücel YH. Human glaucoma and neural degeneration in intracranial optic nerve, lateral geniculate nucleus, and visual cortex. Br J Ophthalmol. 2006;90: 674–678. doi:10.1136/bjo.2005.086769

17. Smith JC, Zhang KY, Sladek A, Thompson J, Bierlein ER, Bhandari A, et al. Loss of Retinogeniculate Synaptic Function in the DBA/2J Mouse Model of Glaucoma. eNeuro. 2022;9: ENEURO.0421-22.2022. doi:10.1523/ENEURO.0421-22.2022

18. Van Hook MJ. Influences of Glaucoma on the Structure and Function of Synapses in the Visual System. Antioxid Redox Signal. 2022;37: 842–861. doi:10.1089/ars.2021.0253

19. Van Hook MJ, Monaco C, Bierlein ER, Smith JC. Neuronal and Synaptic Plasticity in the Visual Thalamus in Mouse Models of Glaucoma. Front Cell Neurosci. 2020;14: 626056. doi:10.3389/fncel.2020.626056

20. Crish SD, Calkins DJ. Central visual pathways in glaucoma: evidence for distal mechanisms of neuronal self-repair. J Neuroophthalmol. 2015;35 Suppl 1: S29–37. doi:10.1097/WNO.0000000000000291

21. Crish SD, Sappington RM, Inman DM, Horner PJ, Calkins DJ. Distal axonopathy with structural persistence in glaucomatous neurodegeneration. Proc Natl Acad Sci USA. 2010;107: 5196–5201. doi:10.1073/pnas.0913141107

22. Bhandari A, Smith JC, Zhang Y, Jensen AA, Reid L, Goeser T, et al. Early-Stage Ocular Hypertension Alters Retinal Ganglion Cell Synaptic Transmission in the Visual Thalamus. Frontiers in Cellular Neuroscience. 2019;13. doi:10.3389/fncel.2019.00426

23. Chen H, Zhao Y, Liu M, Feng L, Puyang Z, Yi J, et al. Progressive degeneration of retinal and superior collicular functions in mice with sustained ocular hypertension. Invest Ophthalmol Vis Sci. 2015;56: 1971–1984. doi:10.1167/iovs.14-15691

24. Smith MA, Xia CZ, Dengler-Crish CM, Fening KM, Inman DM, Schofield BR, et al. Persistence of intact retinal ganglion cell terminals after axonal transport loss in the DBA/2J mouse model of glaucoma. J Comp Neurol. 2016;524: 3503–3517. doi:10.1002/cne.24012

25. Guido W. Development, form, and function of the mouse visual thalamus. J Neurophysiol. 2018;120: 211–225. doi:10.1152/jn.00651.2017

26. Kerschensteiner D, Guido W. Organization of the dorsal lateral geniculate nucleus in the mouse. Vis Neurosci. 2017;34: E008. doi:10.1017/S0952523817000062

27. Schafer DP, Stevens B, Bennett ML, Bennett FC. Role of Microglia in Central Nervous System Development and Plasticity. Cold Spring Harb Perspect Biol. 2024; a041810. doi:10.1101/cshperspect.a041810

28. Gao C, Jiang J, Tan Y, Chen S. Microglia in neurodegenerative diseases: mechanism and potential therapeutic targets. Signal Transduct Target Ther. 2023;8: 359. doi:10.1038/s41392-023-01588-0

29. Hammond TR, Robinton D, Stevens B. Microglia and the Brain: Complementary Partners in Development and Disease. Annu Rev Cell Dev Biol. 2018;34: 523–544. doi:10.1146/annurev-cellbio-100616-060509

30. Salkar A, Wall RV, Basavarajappa D, Chitranshi N, Parilla GE, Mirzaei M, et al. Glial Cell Activation and Immune Responses in Glaucoma: A Systematic Review of Human Postmortem Studies of the Retina and Optic Nerve. Aging Dis. 2024;15: 2069–2083. doi:10.14336/AD.2024.0103

31. Diemler CA, MacLean M, Heuer SE, Hewes AA, Marola OJ, Libby RT, et al. Microglia depletion leads to increased susceptibility to ocular hypertension-dependent glaucoma. Front Aging Neurosci. 2024;16: 1396443. doi:10.3389/fnagi.2024.1396443

32. Bosco A, Steele MR, Vetter ML. Early microglia activation in a mouse model of chronic glaucoma. J Comp Neurol. 2011;519: 599–620. doi:10.1002/cne.22516

33. Bosco A, Inman DM, Steele MR, Wu G, Soto I, Marsh-Armstrong N, et al. Reduced retina microglial activation and improved optic nerve integrity with minocycline treatment in the DBA/2J mouse model of glaucoma. Invest Ophthalmol Vis Sci. 2008;49: 1437–1446. doi:10.1167/iovs.07-1337

34. Bosco A, Breen KT, Anderson SR, Steele MR, Calkins DJ, Vetter ML. Glial coverage in the optic nerve expands in proportion to optic axon loss in chronic mouse glaucoma. Exp Eye Res. 2016;150: 34–43. doi:10.1016/j.exer.2016.01.014

35. Libby RT, Anderson MG, Pang I-H, Robinson ZH, Savinova OV, Cosma IM, et al. Inherited glaucoma in DBA/2J mice: pertinent disease features for studying the neurodegeneration. Vis Neurosci. 2005;22: 637–648. doi:10.1017/S0952523805225130

36. Howell GR, Libby RT, Marchant JK, Wilson LA, Cosma IM, Smith RS, et al. Absence of glaucoma in DBA/2J mice homozygous for wild-type versions of Gpnmb and Tyrp1. BMC Genet. 2007;8: 45. doi:10.1186/1471-2156-8-45

37. Anderson MG, Smith RS, Hawes NL, Zabaleta A, Chang B, Wiggs JL, et al. Mutations in genes encoding melanosomal proteins cause pigmentary glaucoma in DBA/2J mice. Nat Genet. 2002;30: 81–85. doi:10.1038/ng794

38. John SW, Smith RS, Savinova OV, Hawes NL, Chang B, Turnbull D, et al. Essential iris atrophy, pigment dispersion, and glaucoma in DBA/2J mice. Invest Ophthalmol Vis Sci. 1998;39: 951–962.

39. Taylor SE, Morganti-Kossmann C, Lifshitz J, Ziebell JM. Rod microglia: a morphological definition. PLoS One. 2014;9: e97096. doi:10.1371/journal.pone.0097096

40. Holloway OG, Canty AJ, King AE, Ziebell JM. Rod microglia and their role in neurological diseases. Semin Cell Dev Biol. 2019;94: 96–103. doi:10.1016/j.semcdb.2019.02.005

41. Au NPB, Ma CHE. Recent Advances in the Study of Bipolar/Rod-Shaped Microglia and their Roles in Neurodegeneration. Front Aging Neurosci. 2017;9: 128. doi:10.3389/fnagi.2017.00128

42. Giordano KR, Denman CR, Dubisch PS, Akhter M, Lifshitz J. An update on the rod microglia variant in experimental and clinical brain injury and disease. Brain Commun. 2021;3: fcaa227. doi:10.1093/braincomms/fcaa227

43. Stephan AH, Barres BA, Stevens B. The complement system: an unexpected role in synaptic pruning during development and disease. Annu Rev Neurosci. 2012;35: 369–389. doi:10.1146/annurev-neuro-061010-113810

44. Rosen AM, Stevens B. The role of the classical complement cascade in synapse loss during development and glaucoma. Adv Exp Med Biol. 2010;703: 75–93. doi:10.1007/978-1-4419-5635-4_6

45. Young K, Morrison H. Quantifying Microglia Morphology from Photomicrographs of Immunohistochemistry Prepared Tissue Using ImageJ. J Vis Exp. 2018; 57648. doi:10.3791/57648

46. Morrison H, Young K, Qureshi M, Rowe RK, Lifshitz J. Quantitative microglia analyses reveal diverse morphologic responses in the rat cortex after diffuse brain injury. Sci Rep. 2017;7: 13211. doi:10.1038/s41598-017-13581-z

47. Karperien A, Ahammer H, Jelinek HF. Quantitating the subtleties of microglial morphology with fractal analysis. Front Cell Neurosci. 2013;7: 3. doi:10.3389/fncel.2013.00003

48. Stevens B, Allen NJ, Vazquez LE, Howell GR, Christopherson KS, Nouri N, et al. The classical complement cascade mediates CNS synapse elimination. Cell. 2007;131: 1164–1178. doi:10.1016/j.cell.2007.10.036

49. Schafer DP, Lehrman EK, Kautzman AG, Koyama R, Mardinly AR, Yamasaki R, et al. Microglia sculpt postnatal neural circuits in an activity and complement- dependent manner. Neuron. 2012;74: 691–705. doi:10.1016/j.neuron.2012.03.026

50. Reddaway J, Richardson PE, Bevan RJ, Stoneman J, Palombo M. Microglial morphometric analysis: so many options, so little consistency. Front Neuroinform. 2023;17: 1211188. doi:10.3389/fninf.2023.1211188

51. Paolicelli RC, Sierra A, Stevens B, Tremblay M-E, Aguzzi A, Ajami B, et al. Microglia states and nomenclature: A field at its crossroads. Neuron. 2022;110: 3458–3483. doi:10.1016/j.neuron.2022.10.020

52. Ziebell JM, Taylor SE, Cao T, Harrison JL, Lifshitz J. Rod microglia: elongation, alignment, and coupling to form trains across the somatosensory cortex after experimental diffuse brain injury. J Neuroinflammation. 2012;9: 247. doi:10.1186/1742-2094-9-247

53. Tam WY, Ma CHE. Bipolar/rod-shaped microglia are proliferating microglia with distinct M1/M2 phenotypes. Sci Rep. 2014;4: 7279. doi:10.1038/srep07279

54. Witcher KG, Bray CE, Dziabis JE, McKim DB, Benner BN, Rowe RK, et al. Traumatic brain injury-induced neuronal damage in the somatosensory cortex causes formation of rod-shaped microglia that promote astrogliosis and persistent neuroinflammation. Glia. 2018;66: 2719–2736. doi:10.1002/glia.23523

55. Tribble JR, Kokkali E, Otmani A, Plastino F, Lardner E, Vohra R, et al. When Is a Control Not a Control? Reactive Microglia Occur Throughout the Control Contralateral Pathway of Retinal Ganglion Cell Projections in Experimental Glaucoma. Transl Vis Sci Technol. 2021;10: 22. doi:10.1167/tvst.10.1.22

56. Bhandari A, Ward TW, Smith J, Van Hook MJ. Structural and Functional Plasticity in the Dorsolateral Geniculate Nucleus of Mice following Bilateral Enucleation. Neuroscience. 2022;488: 44–59. doi:10.1016/j.neuroscience.2022.01.029

57. Rojas B, Gallego BI, Ramírez AI, Salazar JJ, de Hoz R, Valiente-Soriano FJ, et al. Microglia in mouse retina contralateral to experimental glaucoma exhibit multiple signs of activation in all retinal layers. J Neuroinflammation. 2014;11: 133. doi:10.1186/1742-2094-11-133

58. Yuan T-F, Liang Y-X, Peng B, Lin B, So K-F. Local proliferation is the main source of rod microglia after optic nerve transection. Sci Rep. 2015;5: 10788. doi:10.1038/srep10788

59. de Hoz R, Gallego BI, Ramírez AI, Rojas B, Salazar JJ, Valiente-Soriano FJ, et al. Rod-like microglia are restricted to eyes with laser-induced ocular hypertension but absent from the microglial changes in the contralateral untreated eye. PLoS One. 2013;8: e83733. doi:10.1371/journal.pone.0083733

60. Potru PS, Spittau B. CD74: a prospective marker for reactive microglia? Neural Regen Res. 2023;18: 2673–2674. doi:10.4103/1673-5374.371350

61. Schetters STT, Gomez-Nicola D, Garcia-Vallejo JJ, Van Kooyk Y. Neuroinflammation: Microglia and T Cells Get Ready to Tango. Front Immunol. 2017;8: 1905. doi:10.3389/fimmu.2017.01905

62. Wyss-Coray T, Mucke L. Inflammation in neurodegenerative disease--a double- edged sword. Neuron. 2002;35: 419–432. doi:10.1016/s0896-6273(02)00794-8

63. Neumann H, Boucraut J, Hahnel C, Misgeld T, Wekerle H. Neuronal control of MHC class II inducibility in rat astrocytes and microglia. Eur J Neurosci. 1996;8: 2582–2590. doi:10.1111/j.1460-9568.1996.tb01552.x

64. Wlodarczyk A, Holtman IR, Krueger M, Yogev N, Bruttger J, Khorooshi R, et al. A novel microglial subset plays a key role in myelinogenesis in developing brain. EMBO J. 2017;36: 3292–3308. doi:10.15252/embj.201696056

65. Jia J, Zheng L, Ye L, Chen J, Shu S, Xu S, et al. CD11c+ microglia promote white matter repair after ischemic stroke. Cell Death Dis. 2023;14: 156. doi:10.1038/s41419-023-05689-0

66. Benmamar-Badel A, Owens T, Wlodarczyk A. Protective Microglial Subset in Development, Aging, and Disease: Lessons From Transcriptomic Studies. Front Immunol. 2020;11: 430. doi:10.3389/fimmu.2020.00430

67. Yang R-Y, Rabinovich GA, Liu F-T. Galectins: structure, function and therapeutic potential. Expert Rev Mol Med. 2008;10: e17. doi:10.1017/S1462399408000719

68. Thomas L, Pasquini LA. Galectin-3-Mediated Glial Crosstalk Drives Oligodendrocyte Differentiation and (Re)myelination. Front Cell Neurosci. 2018;12: 297. doi:10.3389/fncel.2018.00297

69. García-Revilla J, Boza-Serrano A, Espinosa-Oliva AM, Soto MS, Deierborg T, Ruiz R, et al. Galectin-3, a rising star in modulating microglia activation under conditions of neurodegeneration. Cell Death Dis. 2022;13: 628. doi:10.1038/s41419-022-05058-3

70. Hoyos HC, Rinaldi M, Mendez-Huergo SP, Marder M, Rabinovich GA, Pasquini JM, et al. Galectin-3 controls the response of microglial cells to limit cuprizone-induced demyelination. Neurobiol Dis. 2014;62: 441–455. doi:10.1016/j.nbd.2013.10.023

71. Elkabes S, DiCicco-Bloom EM, Black IB. Brain microglia/macrophages express neurotrophins that selectively regulate microglial proliferation and function. J Neurosci. 1996;16: 2508–2521. doi:10.1523/JNEUROSCI.16-08-02508.1996

72. Elkabes S, Peng L, Black IB. Lipopolysaccharide differentially regulates microglial trk receptor and neurotrophin expression. J Neurosci Res. 1998;54: 117–122. doi:10.1002/(SICI)1097-4547(19981001)54:1<117::AID-JNR12>3.0.CO;2-4

73. Tabakman R, Lecht S, Sephanova S, Arien-Zakay H, Lazarovici P. Interactions between the cells of the immune and nervous system: neurotrophins as neuroprotection mediators in CNS injury. Prog Brain Res. 2004;146: 387–401. doi:10.1016/s0079-6123(03)46024-x

74. Neumann H, Misgeld T, Matsumuro K, Wekerle H. Neurotrophins inhibit major histocompatibility class II inducibility of microglia: involvement of the p75 neurotrophin receptor. Proc Natl Acad Sci U S A. 1998;95: 5779–5784. doi:10.1073/pnas.95.10.5779

75. Van Hook MJ. Brain-derived neurotrophic factor is a regulator of synaptic transmission in the adult visual thalamus. J Neurophysiol. 2022;128: 1267–1277. doi:10.1152/jn.00540.2021

76. Lessmann V. Neurotrophin-dependent modulation of glutamatergic synaptic transmission in the mammalian CNS. Gen Pharmacol. 1998;31: 667–674. doi:10.1016/s0306-3623(98)00190-6

77. Lessmann V, Heumann R. Modulation of unitary glutamatergic synapses by neurotrophin-4/5 or brain-derived neurotrophic factor in hippocampal microcultures: presynaptic enhancement depends on pre-established paired-pulse facilitation. Neuroscience. 1998;86: 399–413. doi:10.1016/s0306-4522(98)00035-9

78. Hernández-Echeagaray E. Neurotrophin-3 modulates synaptic transmission. Vitam Horm. 2020;114: 71–89. doi:10.1016/bs.vh.2020.04.008

79. Van Hook MJ, McCool S. Enhanced Synaptic Inhibition in the Dorsolateral Geniculate Nucleus in a Mouse Model of Glaucoma. eNeuro. 2024;11: ENEURO.0263-24.2024. doi:10.1523/ENEURO.0263-24.2024

80. Yu A, Tan LX, Lakkaraju A, Santina LD, Ou Y. Microglia target synaptic sites early during excitatory circuit disassembly in neurodegeneration. bioRxiv. 2024; 2024.06.13.598914. doi:10.1101/2024.06.13.598914

